# A simple new alternative to the linear-quadratic model (and where the LQ model comes from)

**DOI:** 10.1101/2024.11.17.624042

**Authors:** L. M. Bilinsky

**Affiliations:** Bilinsky Research

## Abstract

I present a new dose-survival equation for fitting clonogenic assay data collected for irradiated cells, one motivated by the hypothesis that all cellular activities can be partitioned into two states (“state R” and “state Q”) which differ in their sensitivity to low-LET radiation. The LQ model can be derived from it by taking a Taylor expansion. The empirical observation that rapidly proliferating cancer cells have a straighter dose-survival relationship, while slowly proliferating cancer cells and normal cells have a curvier one featuring a shoulder region, is explained in terms of state R and state Q. The new equation (1) provides a convention for classifying cells as radioresistant, (2) provides a means of reducing, or possibly eliminating, cell cycle phase as a variable in treatment outcome, and (3) may enable standardization of the results reported for clonogenic assays. Finally, a novel hypothesis is offered for the “oxygen enhancement ratio” phenomenon.

## 1. Introduction

Radiation is typically administered in a dose range that serves to “sterilize” cancer cells, rather than kill them in the metabolic sense, where “sterilized” means “no longer indefinitely clonogenic,” i.e., able to give rise to an arbitrarily large number of descendants. The clonogenic assay is the standard means of assessing sensitivity to radiation. To carry out this assay, cells in culture are irradiated with dose *D* (in Grays) of ionizing radiation, then seeded on a Petri dish, well spaced from one another. After a duration of time spanning several doubling times, the percentage of cells that went on to produce colonies of 50 or more cells is tallied. After correcting for the plating efficiency, this percentage is referred to as the “surviving fraction” (*SF*) at dose *D*. This procedure is carried out for several values of *D* in the range of interest, and *SF* versus *D* plotted. One often then wishes to fit a mathematical equation to these data, identifying *SF* as a function of *D*, for use in basic research or in planning a course of treatment with fractionated radiotherapy. For over fifty years, the most commonly assumed functional form has been *SF* 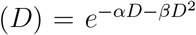. This is known as the “linear-quadratic” (LQ) model. The LQ model has been moderately successful at fitting the clonogenic assay data, but there are data sets for which it fails utterly. See [1] for a discussion of its limitations, which paper also proposes an alternative, known as the power model.

Another set of alternatives is provided by the target models. These assume that a cell contains one or more targets, which must be struck by a photon or particle of radiation one or more times, for the cell to be sterilized. The new model introduced in this paper is of this class. It is a single-target model and I assume that this target is just the genome. However, for the purpose of formulating a dose-survival equation, it is better to think of the target as the entrypoint, upon the cell membrane, of those photons/particles which are ultimately destined for the genome. That is, in the model, the target is the portion of cell membrane occluding the genome. This distinction minimizes confusion later in the paper, where I show that the number of strikes the target must receive for the cell to be sterilized varies with the underlying cellular state, probably due to differences in the resilience of the nuclear envelope.

## 2. Results

In terms of probability theory, the surviving fraction at dose *D* is an estimator of the probability that a cancer cell will *not* be sterilized by this dose of radiation. Our task is thus to determine how this probability varies with *D*. When a cell is irradiated, three questions arise: (1) How many photons/particles of radiation will be emitted by the source in the direction of the cell’s genome? In other words, how many photons/particles will strike the target on the cell membrane? (2) Given that *i* photons/particles strike the target, will they cause genome damage? This requires them to penetrate the nuclear envelope. Let us denote by *u* the probability that genome damage follows *i* strikes upon the target. (3) Given that the target has received *i* strikes and genome damage has occurred, will the cell be sterilized? Repair could take place or the damage, even if unrepaired, might not be such as to cause reproductive death. Let us denote by *p*_0_ the probability that sterilization follows genome damage. With these definitions, we have the following formula for *SF* :

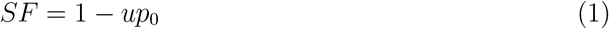

The values of *u* and *p*_0_ in this equation depend upon the exact number of strikes the target receives, the type of radiation being used, and the cancer cell itself.

### 2.1. One strike versus two

Suppose that, for a given population of cancer cells, *u* and *p*_0_ in Eq. 1 do not change with the number of strikes the target receives, provided it receives at least one. The probability of the target receiving at least one strike is *u*_1_ = 1 *−e*^*−kD*^, where *D* is the dose in Grays and *k* is a positive parameter varying directly with the cross-sectional area of the target (equivalently, the genome) and inversely with the per-photon or per-particle energy of the radiation being used. This formula follows from the fact that the number of strikes the target receives is Poisson-distributed with mean *kD*; see *Supplementary Information* for details. *SF* (*D*) for these cells would then be:

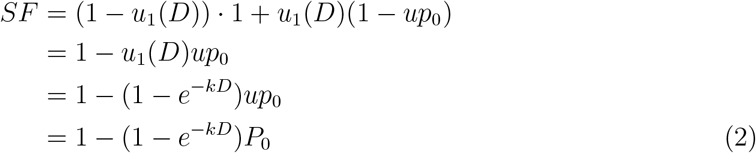

The first line of Eq. 2 follows from the fact that, if the target receives zero strikes of radiation, each cancer cell will go on to produce a colony exceeding 50 cells with probability 1. If one or more strikes are received, sterilization will follow with probability *up*_0_, which product I absorb into the single parameter *P*_0_. A value of *P*_0_ *<* 1 indicates radioresistivity.

Now consider a second population of cancer cells. These cells are never sterilized by a single strike of radiation (*up*_0_ = *P*_0_ = 0 for *i* = 1), but if the target receives two or more strikes (*i ≥* 2), then *u* and *p*_0_ in Eq. 1 are nonzero and constant. The probability that the target will receive at least two strikes is *u*_2_ = 1 *−e*^*−kD*^ *− kDe*^*−kD*^. Hence, *SF* (*D*) for these cells would be:

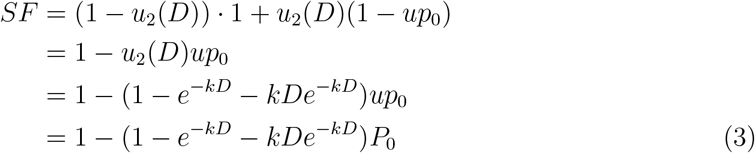

Fig. 1 demonstrates how the shape of the survival curve *SF* (*D*) changes depending upon the number of strikes the target must receive for sterilization to occur.

**Figure 1:**
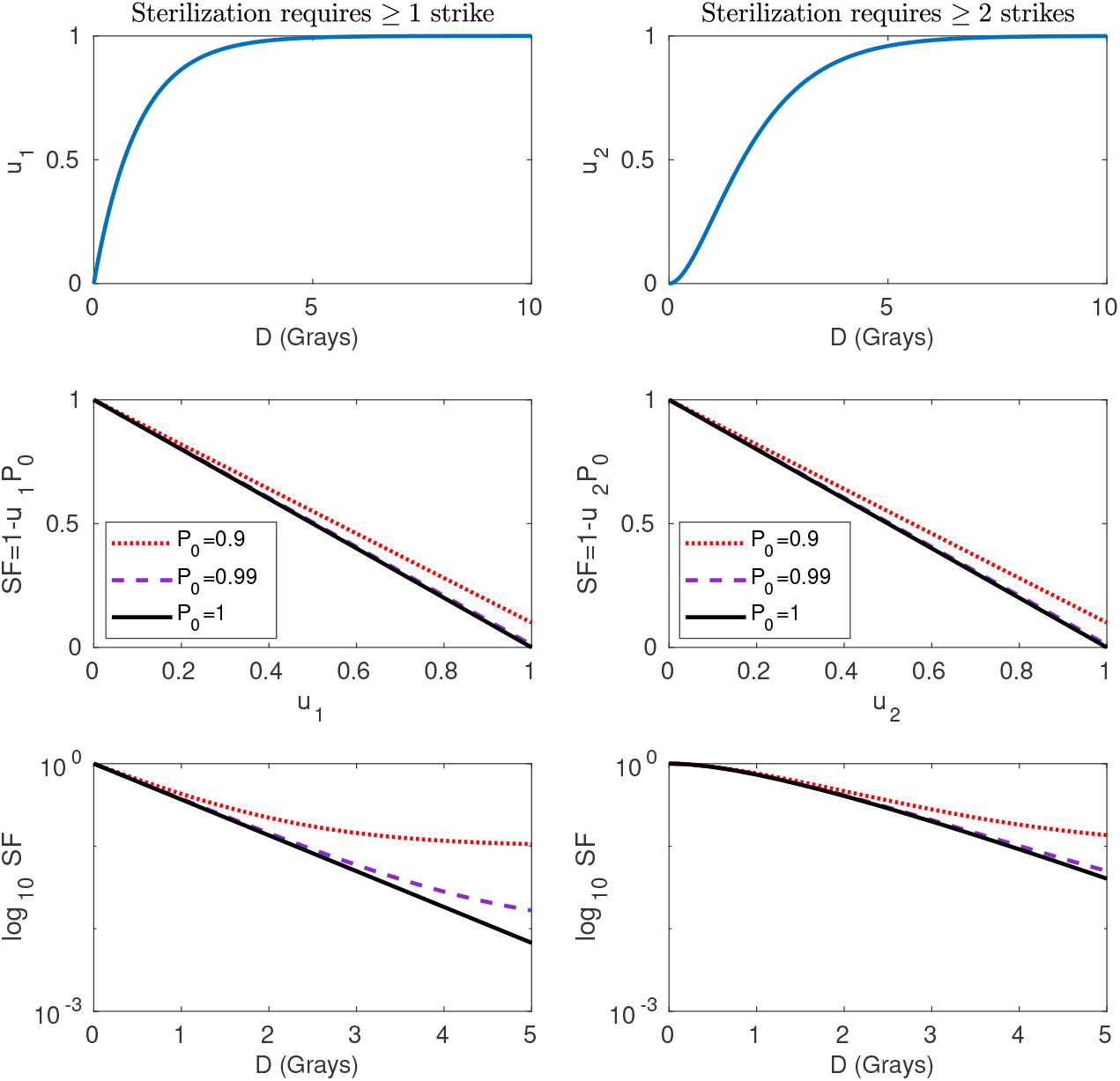
The two typical survival curve shapes obtained using low-LET radiation. *Upper left panel* : *u*_1_ (probability of the target region of the cell membrane receiving one or more strikes of radiation) as a function of *D*. As *D* increases from zero toward infinity, *u*_1_ increases from 0 toward 1 in a way that is determined by Poisson statistics. *Middle left panel* : *SF* as a function of *u*_1_. In the absence of radioresistivity (*P*_0_ = 1), *SF* approaches 0 as *u*_1_ approaches 1. When radioresistivity is present, *SF* approaches 1 *− P*_0_ ≠ 0. *Lower left panel* : *SF* as a function of *D*. In the absence of radioresistivity, *SF* (*D*) plotted on a semilog scale has the shape typically observed for rapidly proliferating cancer cells. When radioresistivity is present, *SF* (*D*) is concave up. *Upper right panel* : *u*_2_ (probability of the target region of the cell membrane receiving two or more strikes of radiation) as a function of *D*. Cf. upper left panel. *Middle right panel* : *SF* as a function of *u*_2_ (identical to middle left panel). *Lower right panel* : The different shape of *u*_2_(*D*) compared with *u*_1_(*D*) results in a different shape for *SF* (*D*). In the absence of radioresistivity, *SF* (*D*) plotted on a semilog scale has the shape typically observed for slowly proliferating cancer cells and normal cells. The shoulder region reflects the fact that two or more strikes have little probability of occurring until *D* is sufficiently large. The shoulder region would have been more prominent had three (or more) strikes been required for sterilization. When radioresistivity is present, *SF* (*D*) eventually becomes concave up as *D* increases. In this figure, *k* = 1.

### 2.2. State R versus state Q hypothesis

Fig. 1 suggests, as a first pass, the following general formula for fitting clonogenic assay data collected using low-LET radiation, for non-radioresistant cells:

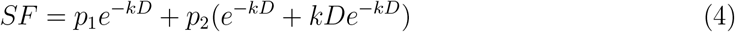

where 0 *≤ p*_1_ *≤* 1, 0 *≤ p*_2_ *≤* 1, and *p*_1_ + *p*_2_ = 1. *p*_1_ (equivalently, *p*_2_) modulates where *SF* (*D*) falls on the spectrum between the two idealized cases shown at the bottom of Fig. 1. Why would clonogenic assay data obey this equation? I suggest the following hypothesis: the cell cycle and *G*_0_ (i.e., all cellular activities) can be partitioned into two phenomenological states. In one of them, a cell will be sterilized if the target region of its cell membrane receives even one strike of low-LET radiation. In the other, the target region must receive two or more strikes for sterilization to occur. *p*_1_ and *p*_2_ in Eq. 4 would then be the fraction of assayed cells in state one and state two, respectively, when radiation is administered.

This is actually a bit of a simplification. Although two strikes, while a cell is in state two, guarantee sterilization for some cell lines (for example, the ovarian cells in Fig. 3), other cell lines are more resilient. The new dose-survival equation, introduced in the next section, takes this into account. State Q is best defined as the complement of state R, the latter being that cellular state in which one strike of low-LET radiation suffices for sterilization.

Decades of clonogenic assay data collected using low-LET radiation indicate that rapidly replicating cancer cells usually have *p*_1_ large (straighter dose-survival curve) while slowly replicating cancer cells and normal, quiescent cells usually have *p*_2_ large (curvier dose-survival curve). In nod to this, I henceforth refer to state one and state two as “state R” and “state Q,” respectively. At the moment of irradiation, *p*_*R*_ of assayed cells will be in state R and *p*_*Q*_ = 1 *− p*_*R*_ will be in state Q. Since replication can be expected to be maximally asynchronous, *p*_*R*_ and *p*_*Q*_ are just the proportions of time cells spend in state R and state Q, respectively:

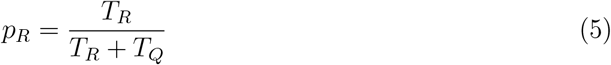

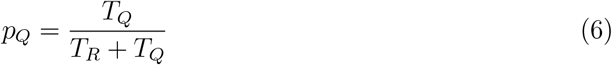

where *T*_*R*_ is the time-duration of state R, *T*_*Q*_ is the time-duration of state Q, and *T*_*D*_ = *T*_*R*_ + *T*_*Q*_ is the doubling time.

The most parsimonious explanation for the observation that *p*_*Q*_ increases as replication rate decreases (doubling time increases) is that the duration of state R varies little across different cancer cell lines, most of the variation in doubling time (17-80 hours [2]) being due to variation in *T*_*Q*_. This motivates a hypothesis, presented later in the paper, for what states R and Q actually represent. For the moment, I emphasize that state R and state Q are not phases of the cell cycle. Rather, each phase of the cell cycle, and *G*_0_, can be partitioned into two sets of activities, those corresponding to state R and those corresponding to state Q. See Fig. 2 for a schematic of the state-R/state-Q hypothesis.

**Figure 2:**
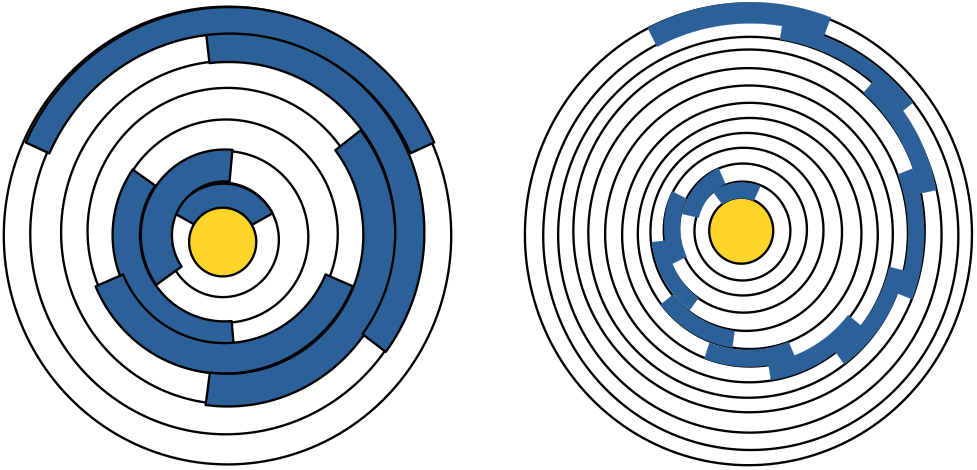
“State Q versus state R” hypothesis. A cancer cell’s progression through the cell cycle can be depicted using a “cancer wheel” diagram. Imagine the cancer wheel as a clock-face with a hand (not shown) which moves clockwise, completing one revolution every *T*_*D*_ hours (the doubling time). Each cancer cell, or set of synchronized cells, occupies one track of the cancer wheel. As the hand moves, cells move with it, progressing through the cell cycle. The dose-survival relationship that has been observed for low-LET radiation can be explained by the hypothesis that the cell cycle can be partitioned into two states, which differ in their sensitivity to low-LET radiation. I call these state R and state Q. When a cell is in state R (blue part of track), it will be sterilized if the target region of the cell membrane receives even one strike of low-LET radiation. When a cell is in state Q (white part of track), more than one strike is needed for this. If a population of cancer cells features maximally asynchronous replication, then all tracks will be equally occupied at the time of irradiation. Hence, 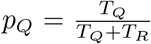 of the cells will be on a white region of track and *p*_*R*_ = 1 *− p*_*Q*_ of the cells will be on a blue region, where *T*_*R*_ is the time-duration of state R, *T*_*Q*_ is the timeduration of state Q, and *T*_*D*_ = *T*_*Q*_ + *T*_*R*_. If *T*_*R*_ is more highly conserved than *T*_*Q*_ across different cell lines, a trend will be observed wherein *p*_*Q*_ is larger for cells with longer doubling times (more slowly replicating cells), causing them to have a curvier dose-survival relationship. *Left panel* : A hypothetical cancer cell line with doubling time 54 hours. 18 of these are spent in state R and 36 are spent in state Q. Hence, *p*_*R*_ = 1*/*3 and *p*_*Q*_ = 2*/*3. *Right panel* : A hypothetical cancer cell line with doubling time 180 hours; this is an extreme example, intended to illustrate the concept. The duration of state R is still 18 hours but the duration of state Q is now 162 hours. Hence, *p*_*R*_ = 0.1 and *p*_*Q*_ = 0.9. In this diagram, each cancer wheel track is partitioned into a single white region and a single blue region. This was done for simplicity. The true pattern is more fragmented, since each phase of the cell cycle is comprised of blue and white regions.

### 2.3. New dose-survival equation

#### 2.3.1. Preliminary remarks regarding dosing

The new dose-survival equation is derived under the assumption that radiation is administered over a fixed time interval Δ*t*, with larger doses being achieved by using a higher dose rate. More typically, the dose rate is held constant and larger doses are achieved by increasing Δ*t*. This may cause the new equation to overestimate *SF* at low doses, as discussed in the *Supplementary Information*.

#### 2.3.2. Equations

Eq. 4 is a special case of the more general Eq. 7 given below, where *p*_1_ and *p*_2_ are now called *p*_*R*_ and *p*_*Q*_, respectively. Eq. 7 is of form *SF* (*D*) = *p*_*R*_*SF*_*R*_(*D*) + *p*_*Q*_*SF*_*Q*_(*D*), where *SF*_*R*_(*D*) and *SF*_*Q*_(*D*) are the theoretical pure-state-R and pure-state-Q dose-survival curves, respectively. The key idea behind this equation is to condition *SF* upon two things: (1) whether a cell is in state R or state Q when radiation is administered and (2) the exact number of strikes the target (i.e., target region of the cell membrane) receives. For each possible choice of (1) and (2) there is a copy of Eq. 1 and a pair of values for *u* and *p*_0_; since these cannot be identified separately from standard clonogenic assay data, they are absorbed into one constant. These are the *P*_0*R,i*_ (*i ≥* 1) and *P*_0*Q,i*_ (*i ≥* 1), which specify the probability of sterilization following *i* strikes upon the target, given that a cell is in state R or state Q. Each sterilization probability reflects all of the possible ways in which *i* strikes could arrive within Δ*t*, the dosing time window. Note that *P*_0*R*,1_ *≤ P*_0*R*,2_ *≤ P*_0*R*,3_ *≤ …*and *P*_0*Q*,1_ *≤ P*_0*Q*,2_ *≤ P*_0*Q*,3_ *≤ …*, since as the number of strikes increases, the probability that sterilization will follow cannot decrease.

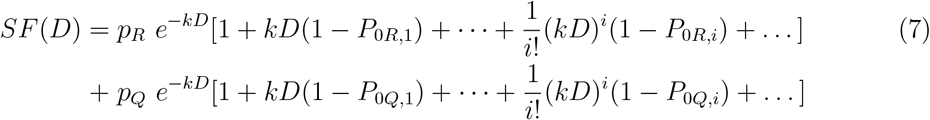

For most cancer cells, we can use Eq. 8 given below. This is a special case of Eq. 7 where we have set *P*_0*R,i*_ = 1 for all *i ≥* 1, *P*_0*Q*,1_ = 0, and *P*_0*Q,i*_ = 1 for all *i ≥* 3. *P*_0*Q*,2_ is a free parameter ranging between 0 and 1. In English this means: if a cell is in state R, then one or more strikes upon the target guarantees sterilization. If a cell is in state Q, then one strike upon the target has no chance of achieving sterilization, two strikes have some chance, and three or more strikes guarantee it.

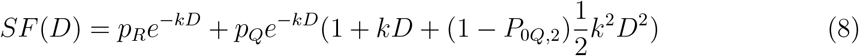

When a good fit to the clonogenic assay data cannot be obtained using Eq. 8, the cells are either ultrasensitive or radioresistant (here I am introducing a convention). Ultrasensitive mutants have *P*_0*Q*,1_ *>* 0 and *P*_0*Q*,2_ = 1, requiring a slight modification of Eq. 8 (not shown, but obvious). Radioresistant cells have either *P*_0*R*,1_ *<* 1 (one strike upon the target does not guarantee sterilization, in state R) and/or *P*_0*Q*,3_ *<* 1 (three strikes upon the target do not guarantee sterilization, in state Q). Eq. 7 must be used for them.

Regarding high-LET radiation, the clonogenic assay data indicate that one strike of this type of radiation guarantees sterilization, whether a cell is in state Q or state R. Since state Q behaves identically to state R, we have simply *SF* = *e*^*−kD*^. However, note that the size of the target region of the cell membrane might be larger for high-LET radiation than for low-LET radiation, resulting in a larger value of *k*, even if the per-photon or per-particle energy were the same. This would occur if such radiation could miss the genome by a wider margin and still be effective.

#### 2.3.3. Terminal linear region

No matter how radioresistant a cell line is, there will be a minimal number *N* such that, if the target receives *N* or more strikes, sterilization is guaranteed. This stems from the fact that no real cell line can withstand an arbitrarily large number of strikes within a finite time interval, unlike the idealized radioresistant cancer cells (*P*_0_ *<* 1) of Fig. 1. In terms of Eq. 7, as *i* increases, the *P*_0*R,i*_ and *P*_0*Q,i*_ eventually attain 1, causing all but a finite number of terms to zero out. Since the critical number of strikes generally differs for state R and state Q, let us call the larger of these *N*. A cell whose target receives *N* or more strikes will be sterilized, regardless of its state. For highly radioresistant cells, like the rr HTB140 cells in Fig. 3, *N* can be quite high; it is 11 for them. A large value of *N* causes the clonogenic assay data to feature a prominent “bustle” region. Beyond the bustle region, the graph of *ln*(*SF* (*D*)) approaches a linear decline with slope *−k*. The terminal linear region constitutes a “signature” assuring us that we have identified the entire graph and can confidently extrapolate out to high doses. If this region cannot be uniquely identified from the clonogenic assay data, then more data must be collected, at higher doses. Note that this region enables us to identify *k* in isolation from other parameters.

**Figure 3:**
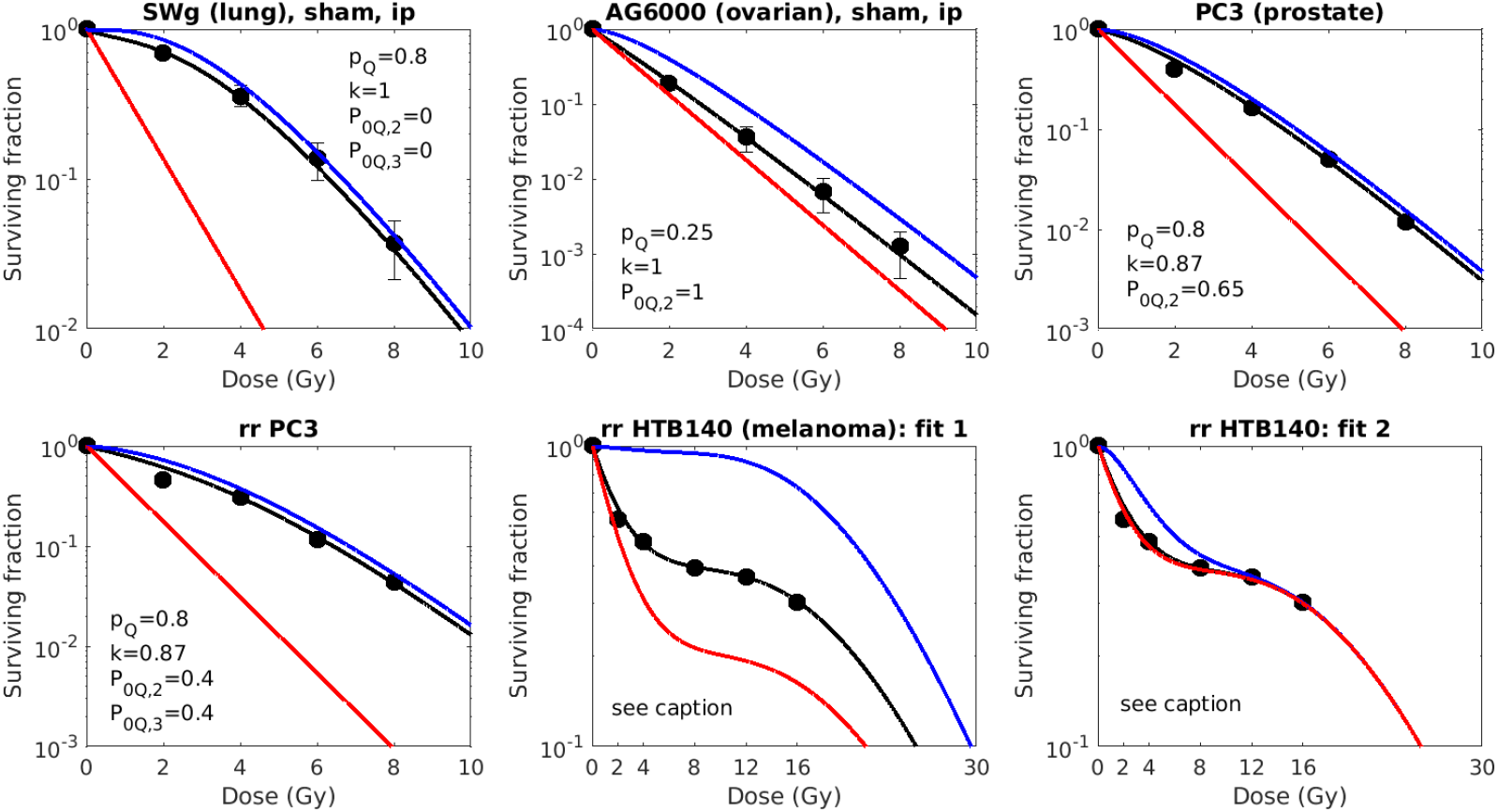
Sample fits of the new dose-survival equation to previously published clonogenic assay data sets. Also shown is the theoretical state-R/state-Q decomposition. *Data sets*: The SWg and AG6000 data sets are reproduced from Fig. 4 of [3]. These data were first published in [4], which paper investigates radiosensitization with gemcitabine. “Sham” indicates that cells were not treated with gemcitabine and “ip” means “immediately plated.” The PC3 and radioresistant PC3 data sets are from Fig. 2 of [5]. Second and third panels of the bottom row present two alternative fits of Eq. 7 to a data set collected for HTB140, an ultra-radioresistant human melanoma cell line. These data are reproduced from [1] and were originally published in [6]. *Model fits*: Optimal fits are shown in black. The short form of the dose-survival equation (Eq. 8) provides a good fit for the AG6000 and PC3 cells. Hence, by the convention introduced in this paper, these two cell lines are normally radiosensitive. The other three cell lines require the full equation (Eq. 7) and are hence classified as radioresistant. Note that *p*_*Q*_ reflects the actual doubling time of irradiated cells, which depends upon the conditions in cell culture. *P*_0*Q*,1_ = 0 for all cell lines, since low-LET radiation is used; aside from this, where *P*_0*Q,i*_ and *P*_0*R,i*_ are not specified, they are equal to 1. *HTB140 cells*: The fit shown in the middle bottom panel has parameter values *p*_*Q*_ = 0.25, *k* = 0.5, *P*_0*R,i*_ = 0.8 (1 *≤ i ≤* 10), *P*_0*Q*,1_ = 0, *P*_0*Q,i*_ = 0.05 (2 *≤ i ≤* 9), *P*_0*Q*,10_ = 0.5. The fit shown in the right bottom panel has parameter values *p*_*Q*_ = 0.1, *k* = 0.5, *P*_0*R,i*_ = 0.62 (1 *≤ i ≤* 9), *P*_0*R*,10_ = 0.72, *P*_0*Q*,1_ = 0, *P*_0*Q,i*_ = 0.62 (2 *≤ i ≤* 9), *P*_0*Q*,10_ = 0.72. *Theoretical decomposition*: The new equation enables us to decompose *SF* (*D*) into theoretical state-R (*SF*_*R*_(*D*), in red) and state-Q (*SF*_*Q*_(*D*), in blue) dose-survival curves. Uses for these are discussed in the *Supplementary Information*.

#### 2.3.4. Sample fits

Fig. 3 shows sample fits of the new dose-survival equation to several published data sets, all of which were collected using low-LET radiation. These fits are discussed in detail in the *Supplementary Information*. Also found there is a discussion of two possible uses for the theoretical state-R and state-Q dose-survival curves. The first is standardization of the results reported for clonogenic assays, which in raw form depend upon the conditions in irradiated cell culture. The second is the minimization, or possible elimination, of the impact of cell cycle phase on the outcome of fractionated radiotherapy.

### 2.4. Derivation of the LQ model

We can write Eq. 7 as follows:

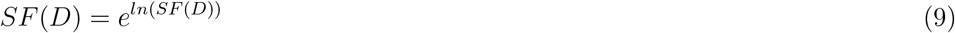

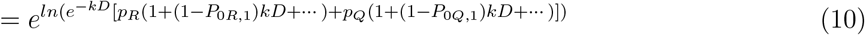

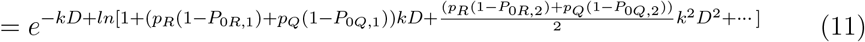

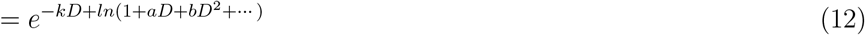

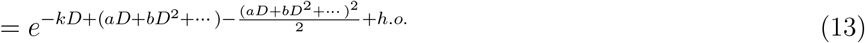

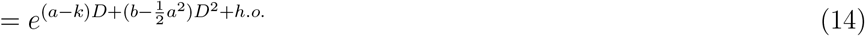

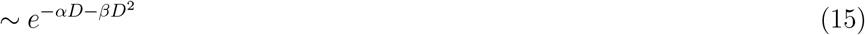

The first line is an identity. In the second line, I insert Eq. 7 for *SF* (*D*). Note that, for any real cell line, only a finite number of terms are represented by the dots; that is, Eq. 7 is in closed form. In the third line, I use the fact that *ln*(*xy*) = *ln*(*x*) + *ln*(*y*), and also rewrite the bracketed expression in a way which makes clear that it is of the form given in line four. In line five, I rewrite line four using the Taylor expansion of *ln*(1 + *x*) about *x* = 0. This gives rise to a power series in *D*. In line six, I collect its like powers. The LQ model (line seven) is obtained by truncating the power series at second order. *α* and *β* are given by:

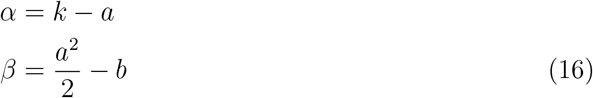

Where

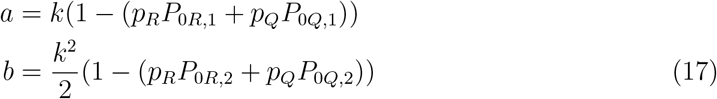

In the *Supplementary Information*, I provide formulas for *α* and *β* in terms of the new model’s parameters for the case where Eq. 8 fits the clonogenic assay data well, and show how *α/β* varies with replication rate. Incidentally, the only time the LQ model is exact is when *SF* = *e*^*−kD*^. In all other cases, the power series in the exponent of *e* has an infinite number of nonzero terms. For example, if *p*_*Q*_ = 1, *P*_0*Q*,1_ = 0, and *P*_0*Q*,2_ = 1, then *SF* = *e*^*−kD*^(1 +*kD*) and *ln* 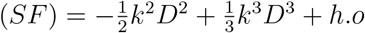.

The LQ model is actually just the second member of an infinite family of models, beginning with *SF* (*D*) = *e*^*−αD*^, the linear model. After the LQ model comes the LQC model, of form 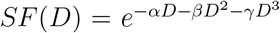. The higher-order members of this family provide an appreciably better fit at high doses, and an appreciably better fit at all doses when radioresistivity is present and *α* and *β* are near zero.

## 3. Discussion

Let us now consider the question of which cellular activities correspond to state R and which to state Q. I speculate that state-R activities are ones for which the rate of ATP hydrolysis within the nucleus is high, and that there is an ANT transporter, analogous to the one located in the inner mitochondrial membrane, embedded in the nuclear envelope. This antiporter acts like a revolving door, exporting one intranuclear ADP to cytoplasm for every cytoplasmic ATP it imports. Such a transporter would be extremely useful, as it would prevent hydrolysis reactions taking place within the nucleus from becoming thermodynamically inhibited as ADP accumulates. Assuming it exists, it would be very active when the rate of ATP hydrolysis within the nucleus is high. This would put a mechanical strain on the nuclear envelope, enabling one particle or photon of low-LET radiation, after entering the cell membrane at the target location, to break the nuclear envelope and still have enough energy left over to inflict damage upon the genome which is irreparable and which renders the cell reproductively unviable. Hence, *P*_0*R*,1_ = 1. Stages of mitosis where no nuclear envelope is present are also subsumed under state R.

State Q is simply the complement of state R, comprised of all cellular activities during which there is not so great a demand for ATP hydrolysis within the nucleus. This probably includes (1) times when the cell is amassing the raw materials (amino acids and lipids) needed to duplicate itself and (2) most of *G*_0_, although note that quiescent cells may still have occasional need for rapid ATP hydrolysis within the nucleus. The reduced mechanical strain on the nuclear envelope during state Q would render it more resilient, so that if a single strike of low-LET radiation were to rupture it, little energy would be left over to damage the enclosed genome, and any damage could be successfully repaired. Hence, *P*_0*Q*,1_ = 0. Two strikes are successful with probability 0 *≤ P*_0*Q*,2_ *≤* 1.

If this hypothesis is correct, it is understandable why state R’s duration would be more highly conserved than state Q’s across different cell lines: a high rate of ATP hydrolysis within the nucleus can only be sustained so long as tasks are being carried out there which require it. These tasks are probably highly conserved across different cell lines and when they are completed, the intense demand for ATP within the nucleus ends. In contrast, amassing the necessary raw materials is an activity which should take an amount of time that is highly variable, if only because different cells have different ratios of surface area to volume. I suspect that this activity accounts for most of the variability, across different cell lines, in *T*_*Q*_. Within a cell line, as nutrients and space in culture become limited, *T*_*R*_ probably doesn’t change, but *T*_*Q*_ will increase.

This line of thought suggests a novel hypothesis for the “oxygen enhancement ratio” (OER) phenomenon. Cancer cells are known for having a much smaller flux through oxidative phosphorylation than normal cells, relying heavily on glycolysis to produce ATP and shunting most of the pyruvate that is produced through lactate dehydrogenase. However, glycolysis itself yields only 2 ATP per glucose, while oxidative phophorylation yields another 30 ATP per glucose [7]. Since the yield is so much higher for oxidative phosphorylation, even if pyruvate enters the Krebs cycle at (for example) 10% the normal rate, oxidative phosophorylation would still be responsible for more than half of the ATP appearing in cytoplasm. The loss of this during hypoxia would cut in half the maximal flux through a nuclear ANT transporter. This would alleviate mechanical strain on the nuclear envelope during state R, decreasing *P*_0*R*,1_ and rendering state R more resistant to low-LET radiation. The same would occur for state Q, if there were a sufficiently high demand for ATP hydrolysis within the nucleus during this state.

It would be informative to treat cells, prior to irradiation, with a (reversible) inhibitor of oxidative phosphorylation and see what effect this has on the dose-survival relationship. If the above hypothesis is correct, *SF* (*D*) will at least become more radiation-resistant. At most, *SF* (*D*) will look as though it were collected during hypoxia. This experiment would decide the extent to which the OER phenomenon is really an OXPHOS phenomenon, mediated by an ANT transporter embedded in the nuclear envelope. A similar hypothesis applies for facultative anaerobes, such as *E. coli*. For them, the role of the nuclear envelope is played by the outer and inner plasma membranes: when oxygen is present and there is a demand for ATP hydrolysis within the bacterium, the F1F0-ATPase embedded in the inner membrane is very active, putting a mechanical strain on the membrane and rendering it more vulnerable to rupture by ionizing radiation.

Even if OXPHOS does not account for most of the OER phenomenon, the latter might still be mediated by the nuclear envelope, rather than by processes downstream of its rupture. Water in the perinuclear space forms hydrogen bonds with head groups on the inner and outer nuclear membranes, stabilizing the nuclear envelope. Liquid water is comprised of water clusters, which are cliques of hydrogen-bonded *H*_2_*O* molecules. Yuan et al. recently reported that *O*_2_ molecules cannot stably exist within water clusters and instead reside at their interfaces [8]. Importantly, they found that as the concentration of dissolved oxygen increases, the average number of water molecules within a cluster decreases. If water cluster size is larger than normal during hypoxia, this may enhance the stability of the nuclear envelope, rendering it more resilient to low-LET radiation. The same effect would occur in the periplasm of bacteria, where water forms hydrogen bonds with head groups on the inner and outer plasma membranes.

## 4. Online Methods

An equation, motivated by first principles, was formulated for the surviving fraction (*SF*) as a function of dose in Grays (*D*). It was then hand-fit to several clonogenic assay data sets drawn from the published literature and the results plotted, using MATLAB R2020b (The MathWorks, Inc., Natick, MA, 2020).

## Author’s contributions

Lydia Bilinsky did all work presented in this paper.

## Acknowledgements

I thank Drs. Trachette Jackson and Patrick Nelson for putting cancer modeling on my radar, while I was an undergraduate student.

## References

[1] H. Li. Invalidity of, and alternative to, the linear-quadratic model as a predictive model for postirradiation cell survival. Cancer Science, 114(7):2931–2938, 2023.

[2] National Cancer Institute. Doubling time of the NCI set of 60 cancerous cell lines (online), 2024.

[3] N. A. P. Franken, A. L. Oei, H. P. Kok, H. M. Rodermond, P. Sminia, J. Crezee, L. J. A. Stalpers, and G. W. Barendsen. Cell survival and radiosensitisation: Modulation of the linear and quadratic parameters of the LQ model (Review). International Journal of Oncology, 42:1501–1515, 2013.

[4] C. van Bree, N. C. Kreder, W. J. P. Loves, N. A. P. Franken, G. J. Peters, and J. Haveman. Sensitivity to ionizing radiation and chemotherapeutic agents in gemcitabine-resistant human tumor cell lines. Int. J. Radiation Oncology Biol. Phys., 54:237–244, 2002.

[5] S. Sideri, F. Petragnano, R. Maggio, S. Petrungaro, A. Catizone, L. Gesualdi, V. De Martino, G. Battafarano, A. Del Fattore, D. Liguoro, P. De Cesaris, A. Filippini, F. Marampon, and A. Riccioli. Radioresistance mechanisms in prostate cancer cell lines surviving ultra-hypo-fractionated EBRT: implications and possible clinical applications. Cancers, 14:5504, 2022.

[6] I. Petrovic, A. Ristic-Fira, D. Todorovic, L. Koricanac, L. Valastro, P. Cirrone, and G. Cuttone. Response of a radioresistant human melanoma cell line along the proton spread-out Bragg peak. International Journal of Radiation Biology, 86(9):742–751, 2010.

[7] W. H. Flurkey. Yield of ATP molecules per glucose molecule (Letter). Journal of Chemical Education, 87(3), 2010.

[8] H. Yuan, Y. Zhang, X. Huang, X. Zhang, J. Li, Y. Huang, K. Li, H. Weng, Y. Xu, and Y. Zhang. Exploration of the existence forms and patterns of dissolved oxygen molecules in water. Nano-Micro Letters, 16, 208, 2024.

