## Supplementary Information for "A simple new alternative to the linear-quadratic model (and where the LQ model comes from)"

---

*Appendix .1. Meaning of the parameter k*

Consider a cancer cell, located a distance $r$ away from a source of ionizing radiation, with no intervening material. Denote as the "target" the region of its cell membrane occluding the genome. Denote by $A_{CS,targ}$ the cross-sectional area presented by the target (equivalently, the genome) to the radiation source. Denote by $A$ the source activity, in mean number of decays per second. Let us administer a single dose of radiation to this cancer cell, by unshielding the source for $\Delta t$ seconds. The number of photons or particles of radiation striking the target during this time interval is Poisson-distributed with mean $\lambda$, where

$$\lambda = A\ \Delta t\ \frac{A_{CS,targ}}{4\pi r^2} \tag{.1}$$

Denote by $A_{CS,cell}$ the cross-sectional area presented by the cell (in its entirety) to the radiation source. The number of photons or particles of radiation striking the cell (anywhere, not just at the target) is Poisson-distributed with mean $\lambda_{cell}$, where

$$\lambda_{cell} = A\ \Delta t\ \frac{A_{CS,cell}}{4\pi r^2} \tag{.2}$$

By definition, the cancer cell receives the following dose in Grays (Joules absorbed energy per kilogram matter):

$$D = \frac{\lambda_{cell} \cdot E}{M_{cell}} \tag{.3}$$

where $E$ is the energy, in Joules, per photon/particle of the type of radiation being used (X-rays, neutrons, $\alpha$-particles, etc.) and $M_{cell}$ is the mass of the cell in kilograms. Note that the actual amount of energy absorbed by the cell is a stochastic random variable, since the number of emitted photons/particles is a stochastic random variable. The dose in Grays is thus the expected value (mean) of the absorbed energy, per kilogram cell.

From Eqs. .2 and .3 we see that when a cancer cell is irradiated, the number of Grays it receives is directly proportional to the ratio $A_{CS,cell}/M_{cell}$. This will be different for cells of different sizes because the ratio of surface area (equal to four times the cross-sectional area) to volume is not constant for a sphere as its radius changes. For example, for the same experimental setup, a larger cell would receive a slightly smaller dose in Grays than would a smaller cell. Even for the same cell, $A_{CS,cell}/M_{cell}$ varies over the cell cycle, since the cell is growing. In light of this, it must be that when a dose is reported in Grays for a clonogenic assay, an average value has been assumed for the ratio $A_{CS,cell}/M_{cell}$. I call this factor $R_{area/mass}$. Combining Eqs. .2 and .3 and substituting $R_{area/mass}$ for $A_{CS,cell}/M_{cell}$, we obtain the following equation for the *reported* dose in Grays:

$$D = A\ \Delta t\ \frac{R_{area/mass}E}{4\pi r^2} \tag{.4}$$

Combining Eqs. .4 and .1 yields:

$$\begin{aligned}\lambda &= D\left(\frac{A_{CS,targ}}{R_{area/mass}E}\right) &(.5)\\ &= kD &(.6)\end{aligned}$$

Hence, $k$ is directly proportional to the cross-sectional area presented by the genome to the radiation source, and inversely proportional to the per-photon/per-particle energy of the type of radiation being used. We can now express, in terms of the *reported* dose in Grays, the probability that the target will receive various numbers of strikes of radiation. The probability that it will receive exactly $i$ strikes is:

$$P_i = \frac{(kD)^i e^{-kD}}{i!} \tag{.7}$$

The probability that it will receive one or more strikes of radiation is:

$$u_1 = 1 - e^{-kD} \tag{.8}$$

The probability that it will receive two or more strikes of radiation is:

$$u_2 = 1 - e^{-kD}(kD + 1) \tag{.9}$$

Fig. .1 shows that as $k$ increases, the graph of $SF$ versus $D$ becomes compressed horizontally, so that smaller doses correspond to a smaller surviving fraction.

*Appendix .2. Considerations that apply when using a constant dose rate*

Eq. 7 assumes that a given number of strikes has a certain probability of causing sterilization, with no other influencing factor (other than whether the cell is in state R or state Q). This is a realistic assumption when radiation is administered over a constant time interval $\Delta t$, the different doses being achieved by varying the dose rate. More typically though, the dose rate is held constant and $\Delta t$ is varied. This poses a potential problem: the probability that sterilization will be achieved by a particular number of strikes should, in general, depend upon the time interval within which they arrive. In particular, it should increase as $\Delta t$ shrinks to zero and the energy of the strikes is deposited at nearly the same time. Since the $P_{0R,i}$ $(i \geq 2)$ and $P_{0Q,i}$ $(i \geq 2)$ vary with $\Delta t$, and $\Delta t$ varies with $D$ (when dose rate is constant), the $P_{0R,i}$ $(i \geq 2)$ and $P_{0Q,i}$ $(i \geq 2)$ are functions of $D$, rather than constant, as Eq. 7 assumes. Despite this, Eq. 7 still fits the clonogenic assay data shown in Fig. 3. Each

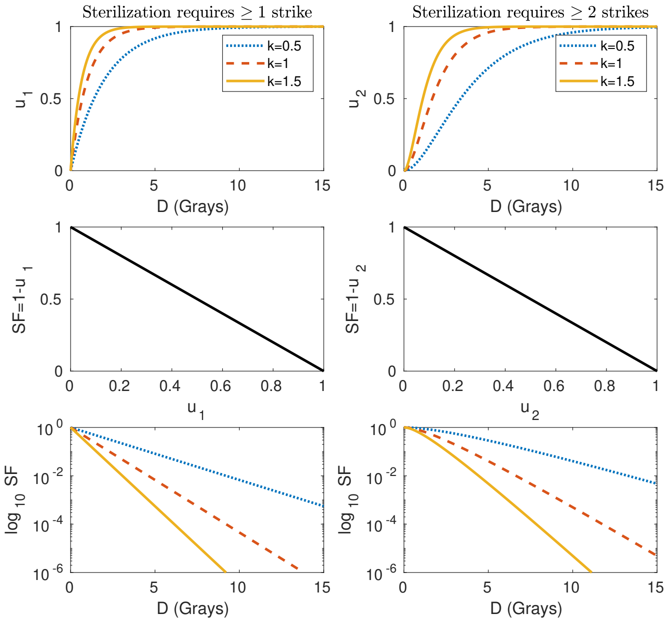


Figure .1: **Effect of varying $k$ on the graph of $SF$ versus $D$.** Compare with Fig. 1. *Upper panels*: The number of strikes received by the target on the cell membrane is Poisson-distributed with mean $kD$. As $D \to \infty$, $u_1$ (the probability of the target receiving at least one strike) and $u_2$ (the probability of the target receiving at least two strikes) approach 1. They do this more rapidly for larger values of $k$. *Middle panels*: These are shown for comparison with Fig. 1. *Bottom panels*: The more rapid approach of $u_1$ and $u_2$ to 1 as $k$ increases translates into a horizontal compression of the graph of $SF$ versus $D$.

of these data sets was collected using a constant dose rate. A possible explanation for this success may be as follows.

The sterilization probabilities $P_{0R,i}$ $(i \geq 2)$ and $P_{0Q,i}$ $(i \geq 2)$ are nonincreasing functions of $\Delta t$ (equivalently, $D$): each should approach a maximal value as $\Delta t \to 0$ (equivalently, $D \to 0$), and as $\Delta t$ (equivalently, $D$) increases, each should decrease or hold steady, eventually plateauing to some constant value. In the plateau region, the time intervals between successive strikes are, with high probability, long enough for the "dust to settle" between strikes. That is, they are long enough that a (damaged) steady state is attained after each strike, so that the exact timing of the next strike is not relevant to the additional damage it effects. In the plateau region, only the number of strikes matter for the sterilization probabilities, not the time gaps between them. Since they are constant in $\Delta t$, they are constant in $D$ and Eq. 7 applies. This analysis assumes that the few minutes over which radiation is administered are not long enough for significant repair of sublethal damage to occur.

The above reasoning suggests that, for the data sets shown in Fig. 3, the time windows over which radiation was delivered were, in the main, long enough for the sterilization probabilities to be in their plateau regions. However, for some data sets, the fit is imperfect at $D = 2$. This indicates that, for this shortest dosing time window, at least one sterilization probability exceeded its plateau-region value. As a general rule, when a constant dose rate is used, the new dose-survival equation is more accurate at higher doses, for which $\Delta t$ is longer; at smaller doses, it may overestimate $SF$. This problem is minimized when using a low dose rate, since the shortest $\Delta t$ will be longer.

Mathematically, it would be much better if radiation were delivered over a constant time interval $\Delta t$, and different doses achieved by varying the dose rate. This would cause Eq. 7 to hold without reservation, improving our ability to fit clonogenic assay data and identify underlying cellular parameters. It would also enable us to use high dose rates without fear of Eq. 7 becoming inaccurate. Higher dose rates are desirable because, for $\Delta t$ small enough, the sterilization probabilities associated with $i \geq 2$ strikes exceed their plateau-region values, allowing us to achieve the same $SF$ using a smaller dose. For example, a high dose rate could be used to shrink the bustle region for highly radioresistant cells.

*Appendix .3. Sample fits*

Fig. 3 (main text) shows sample fits of the new dose-survival equation to several clonogenic assay data sets found in the published literature. The PC3, rr PC3, and HTB140 fits show the predicted imperfection at low doses. The AG6000 fit does not and is not predicted to, since $P_{0Q,2} = 1$: as the time window for dose delivery becomes smaller, a sterilization probability of 1 cannot become any larger. We also do not see the imperfection for the SWg cells. These cells would be classified as highly radioresistant by the convention of this paper. The sterilization probabilities $P_{0Q,2} = 0$ and $P_{0Q,3} = 0$ are compatible with a scenario in which $\Delta t$ (equivalently, $D$) must be very small indeed before $P_{0Q,3}$ and $P_{0Q,2}$ become nonzero. Hence, the imperfection might only appear at, say, $D = 0.5$ Gy.

Let us use Eq. 5 to estimate the duration of state R from the AG6000 data. AG6000 cells are gemcitabine-resistant (i.e., resistant to its action as a chemotherapeutic agent) descendants of A2780 ovarian cancer cells and have an exponential-phase doubling time of 22-24 hours [1]. The data shown were collected for control cells in experiments investigating radiosensitization with gemcitabine. The authors state that the control cells were in the exponential phase of growth when radiation was administered [2]. Since the doubling time is known, we can estimate the duration of state R as $T_R = p_R T_D = 0.75 \cdot 24 = 18$ hours. Let us use this value of $T_R$ to estimate the actual doubling time of the PC3 cells, which were at 80% confluence [3] (i.e., far from the exponential phase) when radiation was administered. We obtain $T_{D,act} = \frac{T_R}{p_R} = \frac{18}{0.2} = 90$ hours. This is very reasonable, given that PC3 cells have a reported exponential-phase doubling time of 33-40 hours [4].

SWg cells are gemcitabine-resistant descendants of SW1573 lung cancer cells; the latter have an exponential-phase doubling time of 22-24 hours [1]. The data shown are for control cells in the same experiments investigating radiosensitization with gemcitabine. Cells were reported to be in the exponential phase of growth when radiation was administered. However, the estimated doubling time for these cells is 90 hours, since $p_R = 0.2$. I suspect that the SWg cells experience contact inhibition, as has been reported for a different lung cancer cell line [5]. Although far from carrying capacity, their actual doubling time, at the moment of irradiation, was probably much longer than 24 hours. The fit shown in Fig. 3 has $p_R$ set to the largest value that is consistent with the data, corresponding to the shortest possible doubling time, under the assumption that state R is normally radiosensitive. If we allow state R to be radioresistant, $p_R$ can have any value. Regardless of what is going on with these cells, they are an exception to the general rule that rapidly replicating cells have a straight dose-survival relationship.

For cells that are extremely radioresistant (ones having a pronounced bustle region), multiple fits are possible. An example of this is provided by the rr HTB140 melanoma cells. The competing fits shown in Fig. 3 differ in how they apportion “responsibility” for the radioresistivity between states Q and R. This translates into different values of $p_Q$ and different theoretical state-R and state-Q dose-survival curves. How could we decide between the two fits? One approach would be to redo the clonogenic assay, this time for cells at a higher confluency (but less than 100%). The true fits, for both confluency conditions, probably share the same $SF_R(D)$, as discussed in the next section. Whether $SF_Q(D)$ is also invariant is less certain.

There is another possible criterion. The HTB140 cells had a doubling time of 24 hours at the time of irradiation [6]. Using this information, the fit shown in the second panel of the bottom row estimates the duration of state R as $T_R = 18$ hours ($T_Q = 6$ hours), in accord with the AG6000 data. The fit shown in the third panel of the bottom row estimates it as $T_R = 21.6$ hours ($T_Q = 2.4$ hours). $T_Q = 2.4$ hours seems unrealistically short (see *Discussion*), indicating that we should go with the fit in the second panel. To establish a rigorous time-duration criterion for deciding between competing fits, $T_R$ and $T_Q$ should be estimated for a large number of normally radiosensitive cells. This would identify (1) the typical range for $T_R$, which should be pretty narrow, and (2) any lower bound on $T_Q$. For this approach to work, we must know cellular doubling times with relative certainty, both for the “database” cell lines used to establish the criterion, and when the criterion is being applied. Toward this end, clonogenic assay data should be collected for cells undergoing exponential growth in medium that is replete with nutrients, so that the doubling time is known with greatest precision.

*Appendix .4. Uses of $SF_R(D)$ and $SF_Q(D)$*

As nutrients and/or space in cell culture become limited, the actual doubling time ($T_{D,act}$) of cells will increase. A hypothesis of this paper is that the duration of state R ($T_R$) will remain constant, assuming that glucose is not so depleted as to limit the rate of ATP synthesis. $T_Q = T_{D,act} - T_R$ should then increase. The question arises: will $SF_Q(D)$ and $SF_R(D)$ change, as the conditions in cell culture change? This matter must be settled empirically, but here I offer some speculation. First, we must discuss the notion of "microstates."

For the sake of simplicity, I have assumed, to this point, that state Q is equally radioresistant (same sterilization probabilities, in response to $i$ strikes upon the target region of the cell membrane) wherever it occurs in the cell cycle. In reality, there may be "shades of white" in Fig. 2, and $SF_Q(D)$, as identified from standard clonogenic assay data, is really an average, having contributions from all the different state-Q "microstates." These microstates differ in the sterilization probabilities assigned to the various numbers of strikes. There would be nothing surprising about the existence of more than one state-Q microstate: the resilience of the nuclear envelope is not the only determinant of sterilization probabilties, which also depend upon the conformation of the genome in space, and, possibly, other factors.

If $SF_Q(D)$ is invariant as the conditions in cell culture change, there is, most likely, only one state-Q microstate. This is a very nice situation, as $SF_Q(D)$ then provides an upper bound on the actual value of $SF$ that a given dose achieves (which is either $SF_R(D)$ or $SF_Q(D)$), no matter where a cancer cell is in its cycle when radiation is administered. Using $SF_Q(D)$, instead of $SF(D)$, to select fraction size would then eliminate cell cycle phase as something which might prevent tumor eradication by treatment's end. Of course, this would come at the cost of administering more radiation.

If $SF_Q(D)$ is not invariant, then there is, most likely, more than one state-Q microstate. As conditions in cell culture change and the actual doubling time increases from its minimal value, reflecting an increase in $T_Q$, it is unlikely that the time durations of the various microstates all increase by the same factor. Since their weights, in determining $SF_Q(D)$, would change, we would observe $SF_Q(D)$ to change. $SF_Q(D)$ is still of use in this case, since it provides an upper bound on the actual $SF$ for those state-Q microstates which are "below average." Using $SF_Q(D)$, rather than $SF(D)$, to select fraction size would at least make less likely a scenario in which an indefinitely clonogenic cancer cell remains at the end of treatment.

If $SF_Q(D)$ does vary with culture conditions, then it is important to use the most radiation-resistant of the $SF_Q(D)$ for this application. That is because some cancer cells within a tumor will have more access to space and nutrients than others, and if we do not use the $SF_Q(D)$ with the smallest sterilization probabilities, there might be some cancer cells for which $SF_Q(D)$ does not satisfy the bounding property just discussed.

Regarding $SF_R(D)$, it is probably invariant with changing conditions in cell culture. Although, for radioresistant cells (more precisely, ones with a radioresistant state R), it is possible for there to be more than one state-R microstate, each microstate's time duration probably does not change as $T_{D,act}$ increases. If each microstate's time duration, divided by $T_R$, remains constant as $T_{D,act}$ increases, then the average $SF_R(D)$ curve will not change.

Another use for $SF_R(D)$ and $SF_Q(D)$ is in standardizing the results reported for clonogenic assays, so that they do not depend upon the conditions in irradiated cell culture. If $SF_R(D)$ and $SF_Q(D)$ are invariant, then they constitute "basis functions" for a given cell line: as conditions in cell culture vary, the value of $p_R$ changes, shifting where $SF(D)$ falls on the spectrum between $SF_R(D)$ and $SF_Q(D)$. $p_R = \frac{T_R}{T_D}$ would be maximal when medium is replete with nutrients and cells are in exponential growth, producing the straightest possible dose-survival relationship for the cell line. In less enriched medium, or at high confluence, the actual doubling time would be much longer than $T_D$, causing $p_R = \frac{T_R}{T_{D,act}}$ to be smaller, and producing a curvier dose-survival relationship. Even if $SF_Q(D)$ is not invariant, this same general picture will hold, and we will be able to report a range for the state-Q sterilization probabilities.

*Appendix .5. $\alpha/\beta$ and replication rate*

For clonogenic assay data sets that are well fit by Eq. 8, we obtain the following formulae for $\alpha$ and $\beta$ in terms of the new model's parameters:

$$\begin{aligned} \alpha &= p_R k \\ \beta &= \frac{k^2}{2}(p_R - 1)(p_R - P_{0Q,2}) \end{aligned} \tag{.10}$$

Viewed as functions of $p_R$, $\alpha$ is a line and $\beta$ is a parabola. Although $\alpha$ is always non-negative, it is possible for $\beta$ to be negative. This occurs, for a given $p_R$, if state Q is resilient enough that $P_{0Q,2} < p_R$. This is consistent with intuition: if state Q were infinitely "shouldery," i.e., totally immune to radiation, so that all state-Q sterilization probabilities (including $P_{0Q,2}$) were zero, $SF(D)$ would simply be $SF(D) = p_Q + p_R e^{-kD}$. The graph of this function is concave up and hence its LQ approximation has $\beta < 0$.

Let us examine how $\alpha$ and $\beta$ change as $p_R$ decreases. As $p_R \to 0$, $\alpha \to 0$ and $\beta \to \frac{k^2}{2} P_{0Q,2}$. Thus, slowly replicating cells have $\alpha/\beta$ small and positive, provided $P_{0Q,2} \neq 0$. If $P_{0Q,2} = 0$, $\alpha/\beta \to -\frac{2}{k}$ as $p_R \to 0$. Assuming the values of $k$ in Fig. 3 are typical, this limit is of order 1. However, this case is not really seen in practice: since both $\alpha$ and $\beta$ are approaching zero, the LQ model would approach simply $SF = 1$, indicating the need for the higher-order terms in $ln(SF(D))$. When confronted with such a data set and armed only with the LQ model, the best-fit values for $\alpha$ and $\beta$ would not be the actual power-series coefficients. For example, the best LQ approximation to Eq. 8 with $k = 1$, $P_{0Q,2} = 0$, and $p_R = 0$, for $0 \leq D \leq 10$, has $\alpha = 0.13$ and $\beta = 0.05$; since these are not the asymptotically correct coefficients, the LQ model does *not* become better and better, for this case, as $D \to 0$ (not shown). This issue exists for any "sufficiently shouldery" data set, such as might be collected for normal, quiescent cells with a particularly resilient state Q.

Let us now see what happens as $p_R$ increases. As $p_R \to P_{0Q,2}$ (the first root of $\beta(p_R)$), $\beta$ approaches zero from the positive direction, causing $\alpha/\beta$ to approach positive infinity. On the other side of this root, $\alpha/\beta$ changes sign. As $p_R$ continues to increase, $\alpha/\beta$ comes in from negative infinity (becomes smaller in magnitude), then shoots out again toward negative infinity as $p_R$ closes in on 1. This assumes that $0 < P_{0Q,2} < 1$. When $P_{0Q,2} = 1$, the analysis is more straightforward: as $p_R$ increases from 0 to 1, $\alpha/\beta$ increases from zero to positive infinity. When $P_{0Q,2} = 0$, then as $p_R$ increases from 0 to 1, $\alpha/\beta$ decreases from $-\frac{2}{k}$ to negative infinity. Thus, we have the general rule that $\alpha/\beta$ is large in magnitude for rapidly replicating cells. Fig. .2 shows plots of $\alpha$, $\beta$, and $\alpha/\beta$ for theoretical cancer cells with $k = 1$ and $P_{0Q,2} = 0.2$.

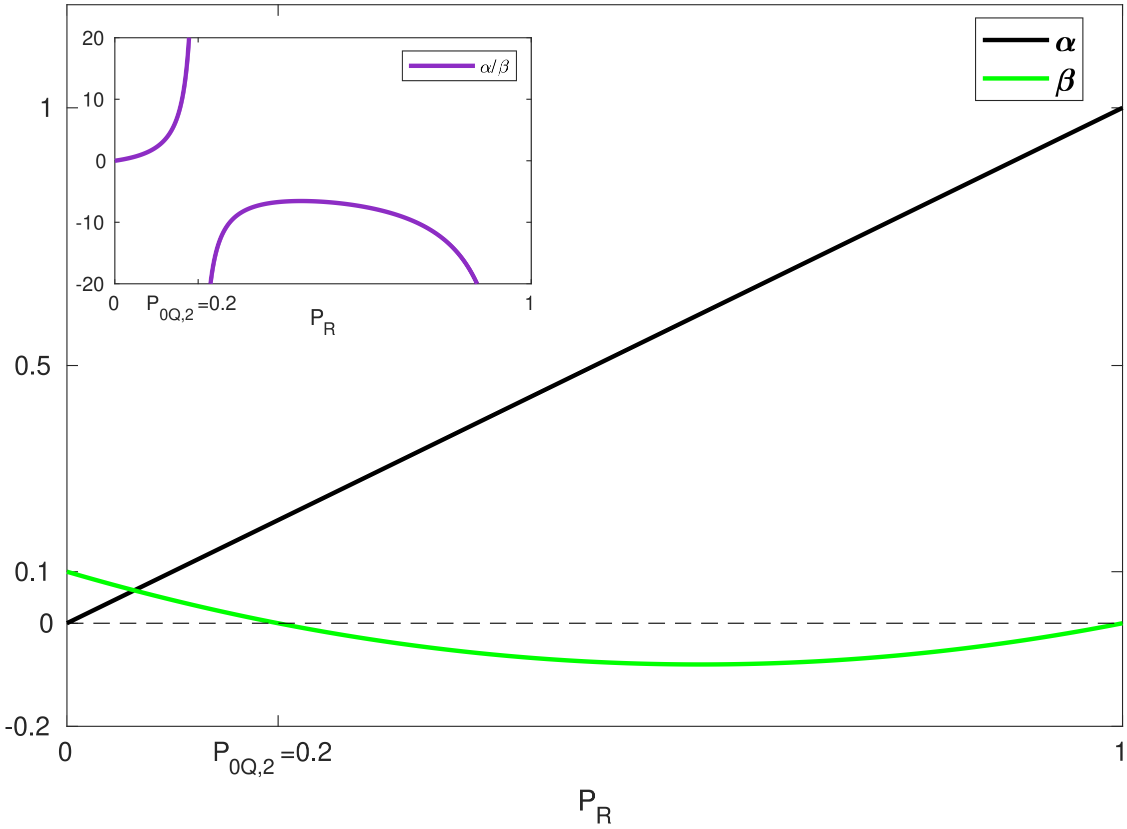


Figure .2: $\alpha$, $\beta$, **and** $\alpha/\beta$ **as functions of** $p_R$**, the proportion of time cells spend in state R.** The LQ model can be obtained from Eq. 7 by taking a Taylor expansion. For cells whose clonogenic assay data are well fit by Eq. 8, $\alpha$ and $\beta$ are as given in Eq. .10. As $p_R \to 0$ (replication rate slows), $\alpha/\beta$ approaches zero from the positive direction. As $p_R \to 1$ (replication rate increases), $|\alpha/\beta|$ approaches infinity. The range of $p_R$ over which $\alpha/\beta$ is positive is $(0, P_{0Q,2})$. In this figure, $k = 1$ and $P_{0Q,2} = 0.2$.

### References


[1] N. A. P. Franken, S. Hovingh, H. Rodermond, L. Stalpers, G. W. Barendsen, and J. Crezee. Radiosensitization with chemotherapeutic agents and hyperthermia: effects on linear-quadratic parameters of radiation cell survival curves. *Journal of Cancer Science and Therapy*, 2011.

[2] C. van Bree, N. C. Kreder, W. J. P. Loves, N. A. P. Franken, G. J. Peters, and J. Haveman. Sensitivity to ionizing radiation and chemotherapeutic agents in gemcitabine-resistant human tumor cell lines. *Int. J. Radiation Oncology Biol. Phys.*, 54:237–244, 2002.

[3] S. Sideri, F. Petragnano, R. Maggio, S. Petrungaro, A. Catizone, L. Gesualdi, V. De Martino, G. Battafarano, A. Del Fattore, D. Liguoro, P. De Cesaris, A. Filippini, F. Marampon, and A. Riccioli. Radioresistance mechanisms in prostate cancer cell lines surviving ultra-hypo-fractionated EBRT: implications and possible clinical applications. *Cancers*, 14:5504, 2022.

[4] C. Su, G. Huang, Y. Chang, Y. Chen, and H. Fang. Analyzing the expression of biomarkers in prostate cancer cell lines. *In Vivo*, 35(3):1545–1548, 2021.

[5] M. Hedman, M. Bergqvist, D. Brattstroem, and O. Brodin. Crowding makes a lung cancer cell line grow slower. *Journal of Thoracic Oncology*, 2(8):S539–S540, 2007. Poster P2-122 presented at 12'th World Conference on Lung Cancer.

[6] I. Petrovic, A. Ristic-Fira, D. Todorovic, L. Koricanac, L. Valastro, P. Cirrone, and G. Cuttone. Response of a radioresistant human melanoma cell line along the proton spread-out Bragg peak. *International Journal of Radiation Biology*, 86(9):742–751, 2010.
